# Omega stabilizes RNA polymerase condensates and contributes to cellular fitness during acid stress

**DOI:** 10.64898/2026.04.03.716306

**Authors:** Stefan Biedzinski, Cyril Haller, Shadi Rajab, Stephanie C. Weber

## Abstract

In eukaryotes, organization of the nucleolus is tightly linked to ribosomal RNA (rRNA) synthesis. External stresses, such as heat shock and acidosis, induce a transition of the nucleolus from a liquid- to solid-like state. Bacterial RNA polymerase (RNAP) condensates share similarities with the nucleolus as they colocalize with pre-rRNA synthesis and their assembly correlates with high rRNA synthesis. In addition, their organization is growth dependent, highlighted by dissolution upon nutritional stress. However, their behavior and biophysical properties during other stresses remain unknown. Here, we find that RNAP condensates persist during acid stress despite an arrest in cell growth. In contrast to fast-growth RNAP condensates, acid-stabilized RNAP condensates become insensitive to drug treatments, suggesting a change in their dynamics. We identify both passive and active mechanisms that contribute to the maintenance of RNAP condensates during acid stress: a drop in intracellular pH and the stringent response. Specifically, the omega subunit of RNAP, which contributes to (p)ppGpp binding site 1, is critical for condensate maintenance during acid stress. In contrast, we show that DksA, a major stress regulator that binds to RNAP, does not contribute to RNAP condensate stabilization. Interestingly, we find that maintenance of RNAP condensates correlates with survival during recovery from acid stress. Our work sheds light on a new aspect of bacterial stress tolerance through regulation of RNAP codensates and a new role for the omega subunit. Our data bring RNAP condensates conceptually one step closer to nucleoli.

## Introduction

**P**hase separation is now recognized as a major mechanism involved in spatial organization in eukaryotic cells (1). For example, the nucleolus is a multi-phase condensate that coordinates the complex process of pre-rRNA synthesis, processing, and ribosomal subunit assembly (2). Recently, similar phenomena have been described in prokaryotes, in which the nucleoid structure is associated with transcription activity through liquid-like organelles, termed RNAP condensates (3, 4). These organelles reversibly assemble in a growth rate-dependent manner, locally concentrating RNAP molecules, which are more mobile than a DNA locus but less mobile than RNAP molecules in the bulk nucleoid (4). These behaviors are characteristic of phase separated liquid-like condensates and differentiate them from aggregates which form irreversibly and in which proteins are immobile (79). RNAP condensates assemble during fast growth in nutrient rich media (5), in which rRNA synthesis accounts for ∼90% of transcription (6). By analogy to the nucleolus, bacterial RNAP condensates were hypothesized to form at sites of rRNA transcription and to facilitate ribosome biogenesis (3, 5). These observations invite further parallels between bacterial RNAP condensates and nucleoli. They could share other similarities, notably regarding their response to stress.

The number and morphology of eukaryotic nucleoli are sensitive to nutrient deprivation, thermal and osmotic stress (7). In addition, the nucleolus transitions from a liquid-to solid-like state during acidosis and heat shock, but not during other stresses such as sodium arsenite or oxidative stress (8, 9). In bacteria, effects of stresses on RNAP condensates remain largely unexplored although the stringent response, a stress response characterized by an increased level of the alarmone (p)ppGpp, is required for the dissolution of RNAP condensates upon amino acid starvation (5). Both nucleoli and RNAP condensates are thus sensitive to stress, yet the response of RNAP condensates to other stresses remains to be explored.

The stringent response has been extensively studied during amino acid starvation (10), in particular using serine hydroxamate (SHX) which inhibits formation of L-Serine tRNA, thus limiting protein synthesis by ribosomes. However, due to the tight link between protein synthesis and cell growth, this experimental approach makes it difficult to isolate the effect of (p)ppGpp alone on RNAP condensation. Increases in cellular (p)ppGpp levels have also been observed during thermal and acidic stress (11, 13, 14). Such stresses are independent of the nutrient supply but their effect on the supramolecular organization of bacterial RNAP has not been addressed to our knowledge.

Bacteria encounter a wide range of external pHs, in the soil, on the skin or in the gastrointestinal tract of their mammalian hosts. Bacteria can use a variety of mechanisms to resist or adapt to acidic environments (15). In mild acidic conditions (5 *<* pH *<* 7), the cell buffers its cytoplasm at a value of 7. However, at extreme external pH values (pH ⩽ 3.5), such as those encountered in the human gastric environment, buffering partially fails and the cytoplasmic pH drops to values between 3.6 and 4.7 (69). This pH drop effectively exposes RNAP condensates to an acidic environment which could perturb molecular conformations and intermolecular interactions, both of which are important for phase separation. Indeed, intracellular acidic environments dissolve liquid-like keratohyalin granules in mouse keratinocytes (16); induce disassembly or assembly of SGs in budding yeast (17); and induce a liquid-to solid-like state transition not only of the nucleolus in human cells (8), but also of stress granules in yeast (17). These observations show that acid stress is a relevant stress to probe the behavior of RNAP condensates during stress regardless of the nutrient supply.

RNAP is comprised of five core subunits (18): beta (β) and beta’ (β’) constitute the enzymatic active core, two alpha (α) subunits interact with gene promoters and transcription factors, and the small omega (ω) subunit is involved in complex assembly and stability. The association of a sigma (σ) factor with the five-subunit core complex forms the holoenzyme and confers promoter specificity (18). RNAP directly binds (p)ppGpp (19) at two different sites: one at the interface between the β’ and ω subunits, close to the enzymatic active center (20, 21); the other at the interface between the β’ subunit and the transcription factor DksA (22). (p)ppGpp alters RNAP activity at specific promoters through conformational changes (23, 24), redirects metabolism (25), and regulates growth rate (26) and thereby participates in stress adaptation.

Here, we explore how bacterial RNAP condensates behave during acid stress, whether the stringent response is involved, and how cellular fitness is affected.

## Results

### RNAP condensates persist during prolonged acid stress

To determine how intracellular pH affects RNAP condensates, we used an *E. coli* strain expressing a fusion protein of mCherry and the β’ subunit (RpoC) from its endogenous locus, as described previously (4). We performed acid stress experiments by diluting a saturated overnight culture into fresh nutrient-rich medium (LB) at 37°C at pH 7.0. Cells were grown for 90 minutes (mid-log phase, t = 0 min) to allow RNAP condensates to form. Hydrochloric acid (HCl) (1 M) was added to lower the medium’s pH to 3.5. A concentrated strong acid was used, rather than a weak acid, to minimize cell culture dilution, as well as to mimic gastric acidity, and to avoid potential anion toxicity of weak acid (27). Optical density (OD) measurements indicate that acid stress arrests cell growth in wild type (WT) cells (Fig. 1A), where growth rate is 0.10 hour^−1^ at pH 3.5 compared to 1.13 hour^−1^ at pH 7.0.

**Fig. 1.**
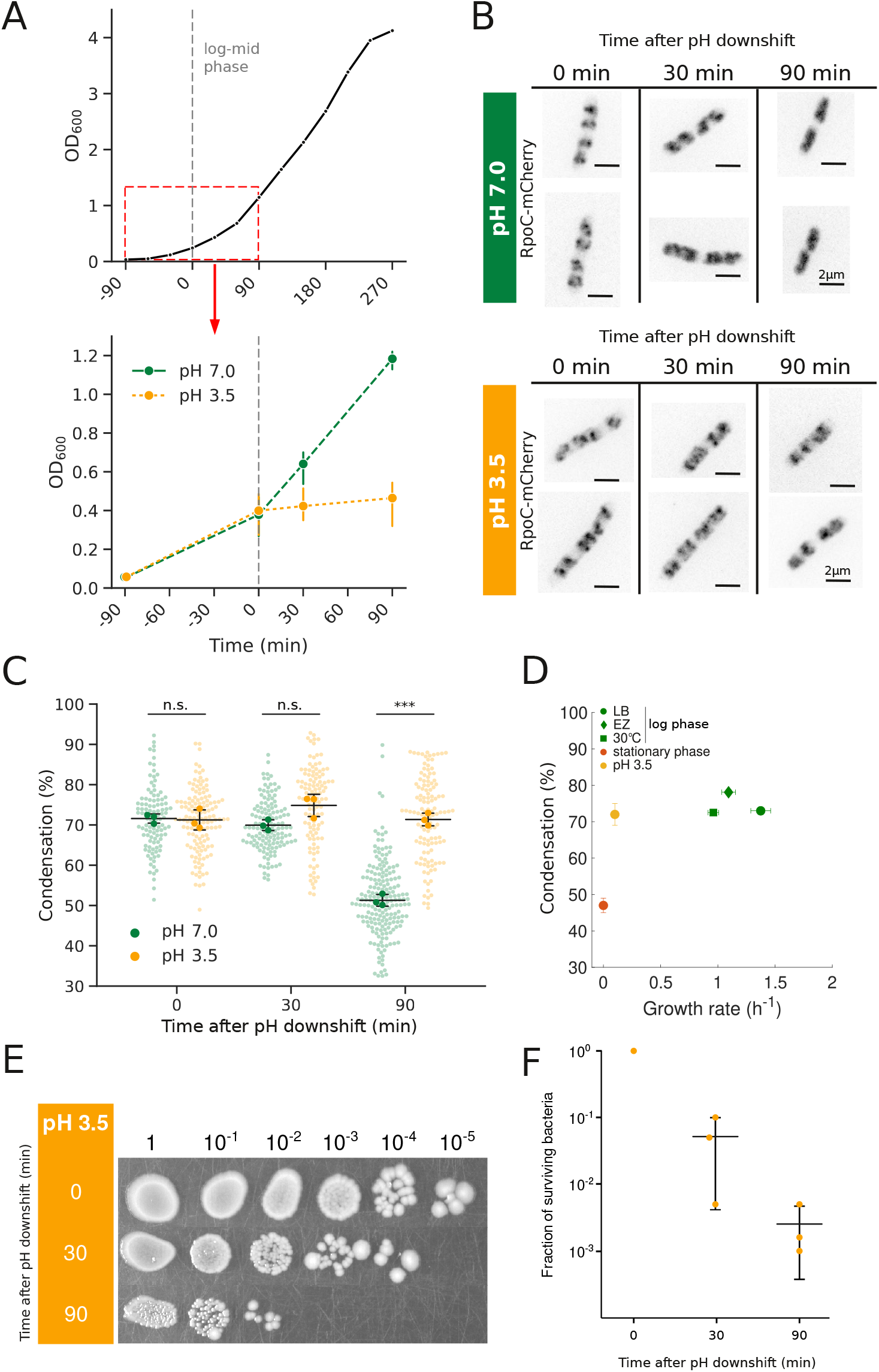
(A) OD_600_ measurements of *E. coli* growing at pH 7 at 37°C in rich media (LB). Red box represents the studied time frame for acid stress. OD_600_ of cultures growing at pH 7.0 (green) or pH 3.5 (yellow) (B) Fluorescence images (contrast inverted) of fixed cells expressing RpoC-mCherry shifted to pH 7.0 or pH 3.5. (C) Condensation of RpoC-mCherry over time after pH downshift (pH7.0 - 0 min: n = 107, 30 min: n = 136, 90 min: n = 191; pH 3.5 - 0 min: n = 135, 30 min: n = 115, 90 min: n = 123). Transparent data points correspond to individual cells; solid markers represent the population mean of each biological replicates; black bars represent mean and standard error of replicates of N = 3 replicates. p-values calculated by t-test. n.s.: non-significant; ***: p < 0.001 (D) Condensation of RNA polymerase in different growth media as a function of growth rate. “EZ” stands for EZ-Rich Defined Medium; “30°C” refers to cells grown in LB at 30°; all other conditions were grown at 37°C. EZ, 30°C, and stationary phase data are taken from (4) (E) Survival assay at various times after pH downshift. Each column represents a dilution of 5μL droplet. Experiment was repeated 3 times. (F) Colony counts of panel E. Values are presented as a fraction relative to t = 0 minute.

During outgrowth, samples were taken at various time points (t = 0 min, t = 30 min, t = 90 min) and fixed with formaldehyde before imaging. At neutral pH, cells contain RNAP condensates during steady-state growth (t = 0, 30 min) (5) which later disappear (t = 90 min) as growth rate decreases (Fig. 1A, B, S1A) (4). The dissolution of condensates correlates with a decrease in cell length (Fig. S1B, C), consistent with their growth rate-dependent nature. When exposed to pH 3.5, cells maintain bright RNAP condensates (Fig. 1B, C) for up to 90 min. Cell length also remains constant during acid stress (Fig. S1C), indicating that RNAP condensates persist despite growth arrest.

To quantify condensation, we used a clustering metric developed in (4) (see Materials and Methods). In *E. coli* cultured at pH 7.0, quantification confirms that condensation decreases from 71 ± 8 % (mean ± standard deviation) at t = 0 min to 52 ± 9 % at t = 90 min (Fig. 1C). In contrast, cells exposed to acidic pH exhibit condensation of 71.2 ± 2 % at t = 0 min, which they maintain to a value of 69 *±* 11 % at t = 90 min (Fig. 1C). To determine whether the difference in condensation between control and acid-stressed cells at 90 min is due to changes in RpoC abundance, we measured total fluorescence intensity per cell across our conditions. While RpoC abundance is lower at pH 7.0 than pH 3.5, the decrease in total fluorescence at pH 7.0 precedes the dissolution of RNAP condensates (Fig. S2A,B,C), suggesting that differences in condensation are not due to changes in protein abundance.

In order to assess cell viability during extreme acid stress, we performed spot assays. We found that cells exposed to pH 3.5 for 30 min form 20 ± 21 times fewer colonies than cells prior to acid stress, and after 90 min they form 400 ± 500 fewer colonies (Fig. 1E, F). Control cells maintained at pH 7.0 showed an increase in viability (Fig. S1D). These results indicate that most of our observations are made on dead cells. Nevertheless, the distribution of our condensation metric at pH 3.5 does not change over time (ANOVA p-value = 0.18) despite the progressive decrease in cell viability. This suggests that RNAP condensate behavior during acid stress is independent of the living state of the bacteria and may instead depend on the physico-chemical properties of the surrounding environment of the condensates.

We also sought to determine the contribution of nucleoid structure to RNAP condensates maintenance. To do so we acid stressed bacteria expressing HupA-mCherry and observed a slight increase of compaction of the nucleoid structure (Fig. S3A, B) suggesting that the underlying structure of the nucleoid could still be contributing to RNAP maintenance.

Previous observations found a tight link between RNAP condensation and growth rate: RNAP condensates are only present in fast-growing log phase cells and dissolve in slow-growing or stationary phase cells (4, 5) (Fig. 1D). Our results show that during acid stress, condensates are still present despite growth arrest and cell death, illustrating true decorrelation between growth and RNAP condensation. This decoupling raises questions as to the driving forces of such condensation, attributed so far only to transcriptional activity (5) and nucleoid structure (3). We next aimed to explore how growth-independent condensates at pH 3.5 behave in conditions known to disrupt RNAP condensates at neutral pH.

### RNAP condensates are insensitive to hexanediol during acid stress

1,6-hexanediol has been used extensively to dissolve liquid-like condensates through disruption of weak hydrophobic interactions (31). Indeed, we previously showed that RNAP condensates are sensitive to hexanediol, with the percentage of condensation decreasing in a reversible and dose-dependent manner (4). However, hexanediol has also been shown to inhibit enzyme activity *in vitro* (80) and to perturb chromatin dynamics and cytoskeletal structure in eukaryotic cells (81, 82) so it should be used with caution. Since cell viability is already low, we applied hexanediol to acid-stressed cells to alter the physicochemical environment surrounding these RNAP condensates and see how they respond. In control cells at pH 7.0, condensation decreases from 72.7 ± 2% to 52.2 ± 2% after 5 minutes of 5% hexanediol exposure (Fig. 2A,B), as observed previously (4). However, at pH 3.5, hexanediol treatment has little or no effect on RNAP condensates, with 64.7 ± 7% condensation before and 61.4 ± 2% after hex treatment (Fig. 2A,B). This result suggests that acid stress stabilizes RNAP condensates, rendering them insensitive to growth arrest and hexanediol. This stabilization could arise through passive, physicochemical effects on RNAP condensates; active cellular responses before death; or a combination of both. *E. coli* can buffer its cytoplasm, maintaining a pH of 7, over a moderate range of environmental conditions (pH 4-8). However, when the environmental pH drops below 4, the cell’s buffering capacity is exceeded, causing a downshift in intracellular pH, which can affect protein solubility and their ability to phase separate (8, 32, 33, 34, 35). In this regard it is necessary to understand how pH relates to RNAP condensates during extreme but also mild acidic stress.

**Fig. 2.**
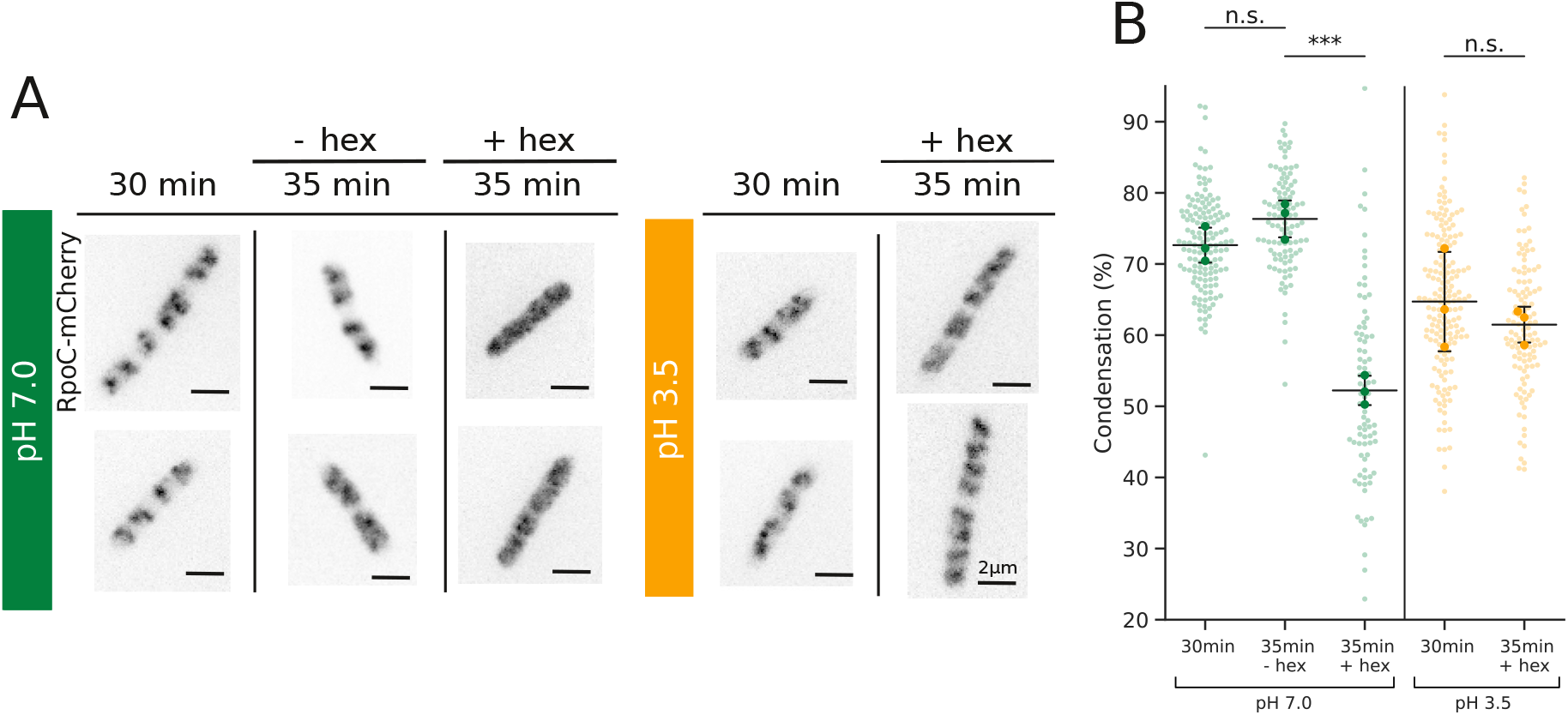
(A) Fluorescence images (contrast inverted) of fixed cells expressing RpoC-mCherry shifted to pH 7.0 or pH 3.5 for 30 min before and after 5 min treatment with rifampicin (rif). (B) Condensation of RpoC-mCherry after pH downshift and treatment with rifampicin. (pH 7.0 - 30 min: n = 93; 35 min - rif: n = 208; 35 min + rif: n = 80; pH 3.5 - 30 min : n = 127; 35 min + rif: n = 145). (C) Fluorescence images (contrast inverted) of fixed cells expressing RpoC-mCherry shifted to pH 7.0 or pH 3.5 for 30 min before and after 5 min treatment with hexanediol (hex). (D) Condensation of RpoC-mCherry after pH downshift and treatment with hexanediol. (pH 7.0 - 30 min : n = 142; 35 min - hex: n = 208; 35 min + hex: n = 84; pH 3.5 - 30 min : n = 173; 35 min + hex: n = 116). Transparent data points correspond to individual cells; solid markers represent the population mean of each biological replicates; black bars represent mean and standard error of N = 3 replicates. p-values calculated by t-test. n.s.: non-significant; **: p < 0.01; ***: p < 0.001

### Intracellular pH downshift correlates with stabilization of RNAP condensates

To confirm pH downshift in our experimental setup, we monitored intracellular pH using SEpHluorin (36), a green fluorescent protein whose fluorescence intensity proportionally decreases with decreasing pH. We observed that extreme acid stress (pH 3.5) lowers intracellular pH to values below 4.0, confirming that intracellular pH becomes close to extracellular pH (Fig. 3A, C). In contrast, exposure to a mild acidic medium (pH 5.5) does not affect intracellular pH, which remains stable at 6.90 ± 0.35 (Fig. 3B, C). This result indicates that intracellular pH buffering remains functional at external pH 5.5 but fails at external pH 3.5, as expected (15).

**Fig. 3.**
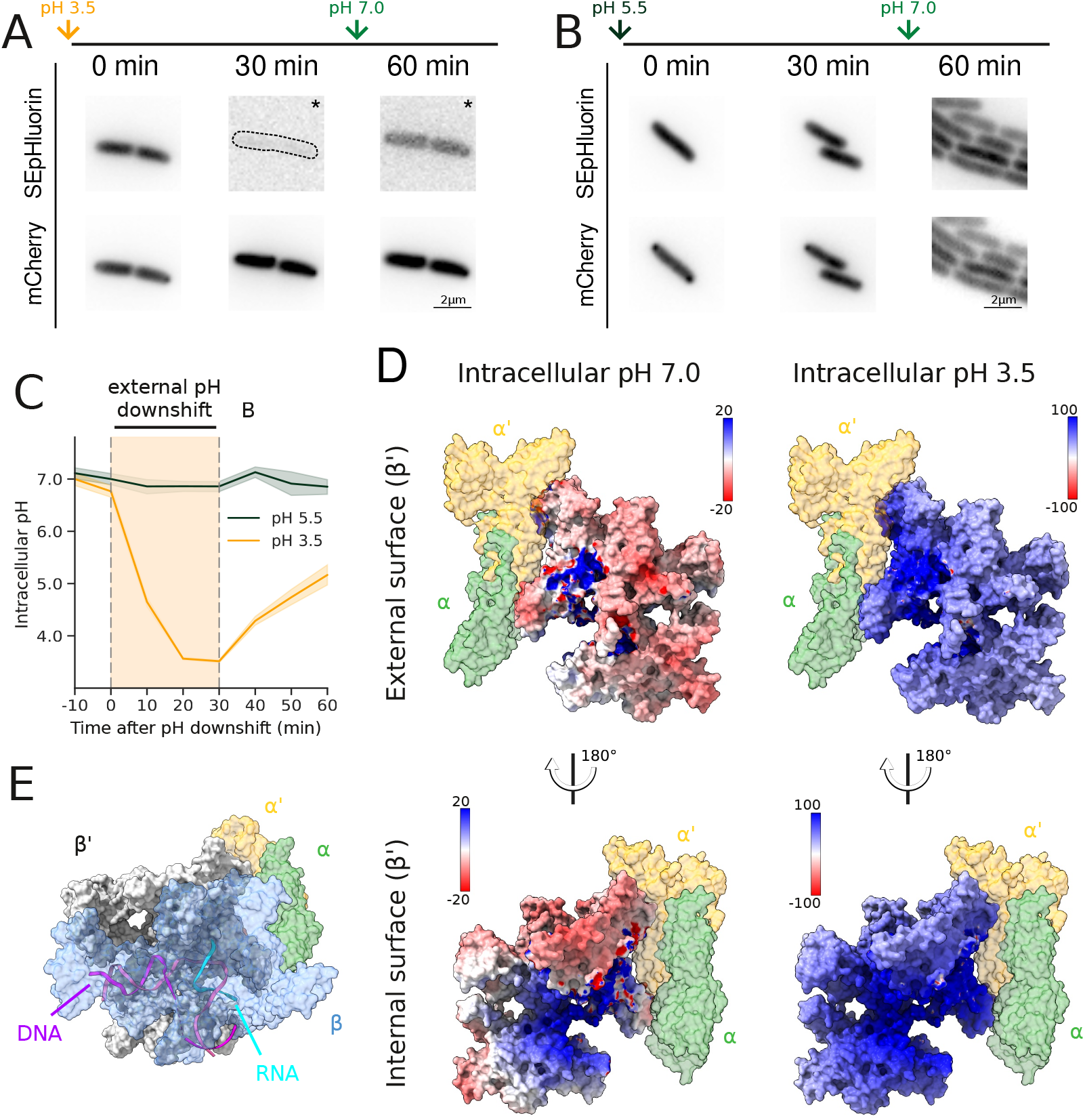
(A) Fluorescence images (contrast inverted) of live cells expressing SEpHluorin-mCherry before, during and after downshift to pH 3.5 or (B) pH 5.5. “*” indicates enhanced contrast for visualization. (C) Fluorescence intensity of SEpHluorin in live cells normalized to fluorescence intensity of mCherry before, during, and after external pH downshift to pH 5.5 or 3.5. Medium was changed from pH 7.0 to pH 5.5 or 3.5 after 10 minutes, and back to pH 7.0 after 30 minutes of stress (yellow region). Thick line represents the mean, shaded region represents standard deviation (SD) across replicates. (D) Molecular structure of the RNAP complex transcribing DNA (PDB #7MKO (79)). The complex is oriented such that the internal surface of the β’ subunit in contact with the DNA template and nascent RNA visible. The ω subunit is located behind β’ and is not visible. (E) Electrostatic potential of the RNAP β’ subunit (PDB #5VSW (76)) (see Materials and Methods for calculation). α (green) and α’ (yellow) subunits are included as spatial references. Electrostatic potential is displayed in kcal/(mol·e) at 298 K, with positive values in blue, negative in red, neutral in white.

Intracellular pH directly impacts the protonation state of amino acids, and thus, a protein’s net charge. Theoretical net charge of an unfolded protein can be calculated with the Henderson–Hasselbalch equation applied to each amino-acid side chain to obtain the proportion of protonated and non-protonated species (see Materials and Methods). Using this method, we calculated the net charge of each subunit of the RNAP core complex at intracellular pH 7.0 and intracellular pH 4.0 (Table 1). Both β and β’ drastically increase in net charge, from -39.3 to +128.1 and from +0.8 to +152.9, respectively. In contrast, ω’s net charge is only modestly affected by acidic pH values (see Table 1), shifting from -4.1 to +8.3. These observations suggest that the two largest subunits of RNAP may be particularly affected by a downshift in cytoplasmic pH.

**Table 1.**
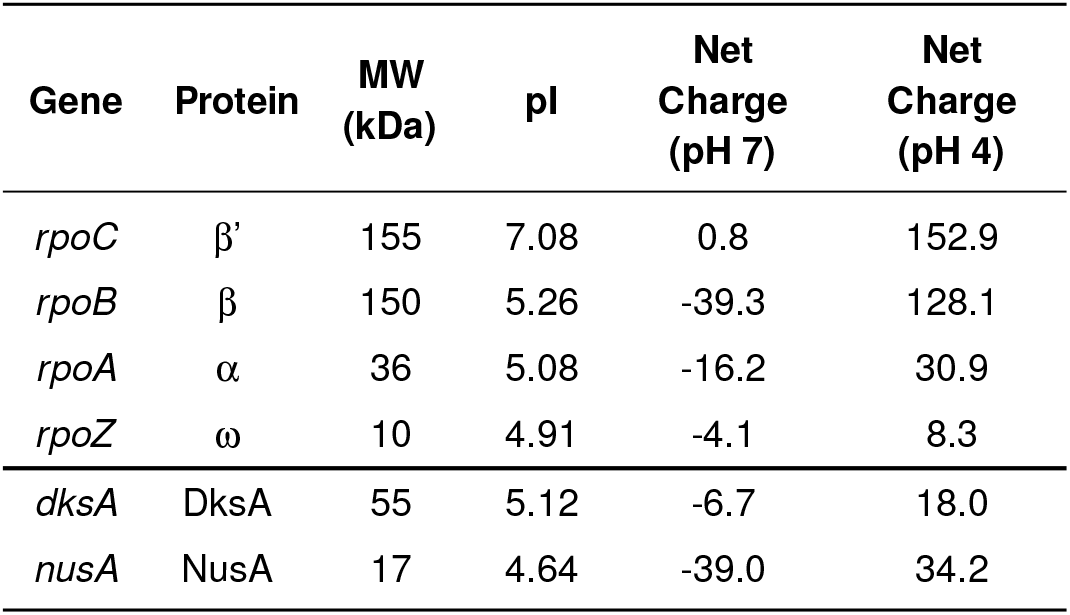
Electrostatic properties of RNAP subunits and associated proteins.

| Gene | Protein | MW<br>(kDa) | pI | Net<br>Charge<br>(pH 7) | Net<br>Charge<br>(pH 4) |
| --- | --- | --- | --- | --- | --- |
| *rpoC* | $\beta'$ | 155 | 7.08 | 0.8 | 152.9 |
| *rpoB* | $\beta$ | 150 | 5.26 | -39.3 | 128.1 |
| *rpoA* | $\alpha$ | 36 | 5.08 | -16.2 | 30.9 |
| *rpoZ* | $\omega$ | 10 | 4.91 | -4.1 | 8.3 |
| *dksA* | DksA | 55 | 5.12 | -6.7 | 18.0 |
| *nusA* | NusA | 17 | 4.64 | -39.0 | 34.2 |

*In vitro* and theoretical studies (33, 35) demonstrate that homotypic protein phase separation is favored at pH values close to the isoelectric point (pI), i.e. when the protein’s net charge is close to 0, and that deviation from the pI suppresses phase separation. In addition, it has been shown that at low pH, histidine protonation alone can be unfavorable to phase separation (37). The RNAP core complex contains in total 56 histidines (β: 21, β’: 19, α: 8). Altogether, these literature results would predict that RNAP condensates should dissolve during acidic stress in contrast to the stabilization that we observe (Fig. 1, 2).

To address this apparent discrepancy, we next considered other components of RNAP condensates. For example, the anti-termination factor NusA is known to colocalize with RNAP (73) and can nucleate droplets *in vitro* and *in vivo* (4). The net charge of NusA shifts from -39.0 at intracellular pH 7.0 to +34.0 at pH 4.0, which could significantly affect the strength of its intermolecular interactions. In addition to proteins, RNAP condensates also contain both DNA and RNA, which have negatively-charged backbones even at pH 4.0.

We next calculated the electrostatic potential of the RNAP core complex. As expected from the increase in net charge, we found a drastic shift from a mostly neutral to a strong positively charged surface of β’ when intracellular pH is shifted from pH 7.0 to pH 4.0 (Fig. 3D), notably on its inner surface, where it interacts with nucleic acids (Fig. 3E). This increase in electrostatic potential on the internal surface would increase the binding strength between RNAP and DNA and RNA, and may explain, at least partially, the increased stability of RNAP condensates observed in our experiments.

In addition to this passive physicochemical mechanism, bacteria have an active pathway to adapt to various stresses, the stringent response (11). This pathway acts directly on RNAP during amino acid starvation, so we next examined whether it contributes to the stabilization of RNAP condensates during acid stress.

### The stringent response is required for acid-induced stabilization of RNAP condensates

In the stringent response, (p)ppGpp is synthesized primarily by RelA, a ribosome-associated protein activated by uncharged tRNAs in response to amino acid starvation, as well as by SpoT, a (p)ppGpp synthase and hydrolase (Fig. 4A). (p)ppGpp binds two different sites on the RNAP complex (22): at the interface between the β’ and ω subunits (site 1) and at the interface of β’ and the DksA transcription factor (site 2) (Fig. 4B). To assess whether the stringent response contributes to stabilization of RNAP condensates during acid stress, we analyzed the phenotypes of 3 different mutants: Δ*relA*, which is partially deficient for (p)ppGpp synthesis; Δ*rpoZ*, which is missing the ω subunit; and finally Δ*dksA*, which lacks the transcription factor DksA.

**Fig. 4.**
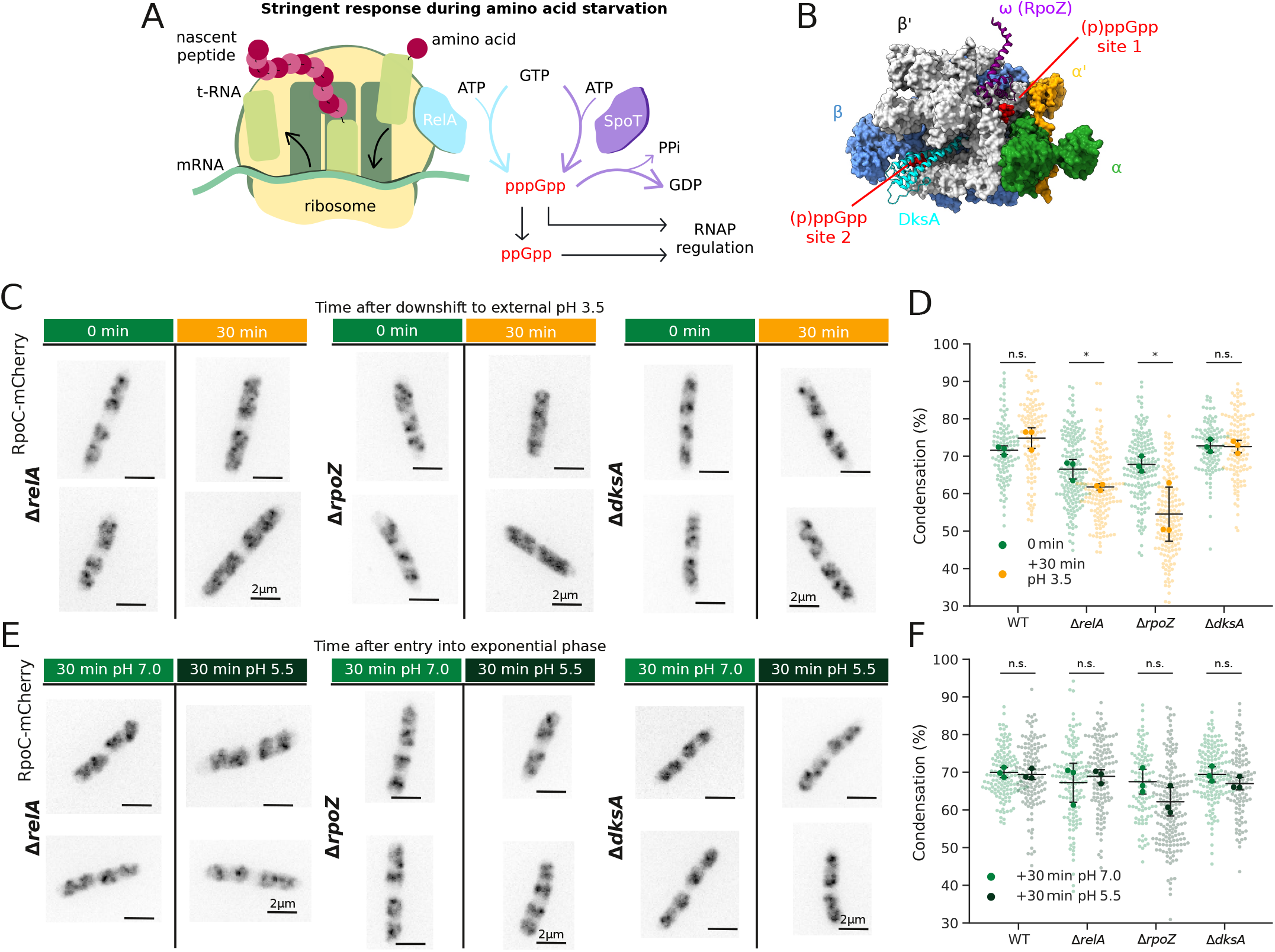
(A) Schematic of the (p)ppGpp synthesis pathway. (B) Molecular 5 core RNAP subunits (PDB#5VSW (**?**)), with DksA and two molecules of (p)ppGpp molecules bound. (C) Fluorescence images (contrast inverted) of fixed Δ*relA*, Δ*rpoZ* and Δ*dksA* cells expressing RpoC-mCherry before (0 min) or after (30 min) downshift to external pH 3.5. (D) Condensation of RpoC-mCherry before or after pH downshift to pH 3.5 of WT (0 min: n=107, 30 min: n=115), Δ*relA* (0 min: n=171, 30 min: n=149), Δ*rpoZ* (0 min: n=164, 30 min: n=166) and Δ*dksA* (0 min: n=103, 30min: n=121). Transparent data points correspond to individual cells; solid markers represent the population mean of each biological replicates; black bars represent mean and standard error of N = 3 replicates. WT data are the same as in Fig. 1C, for comparison. (E) Fluorescence images (contrast inverted) of fixed Δ*relA*, Δ*rpoZ* and Δ*dksA* cells expressing RpoC-mCherry 30 min after entry into exponential phase at pH 7.0 or pH downshift to pH 5.5. (F) Condensation of RpoC-mCherry after 30 min of pH downshift to pH 7.0 and pH 5.5 of WT (pH 7.0: n=136, pH 5.5: n=129), Δ*relA* (pH 7.0: n=137, pH 5.5: n=130), Δ*rpoZ* (pH 7.0 min: n=98, pH 5.5: n=190) and Δ*dksA* (pH 7.0: n=148, pH 5.5: n=110). Transparent data points correspond to individual cells; solid markers represent the population mean of each biological replicates; black bars represent mean and standard error of N = 3 replicates. p-values calculated by ANOVA. n.s.: non-significant; *: p < 0.05

First, we confirmed that cells lacking the dominant (p)ppGpp synthetase RelA (Δ*relA*) assemble RNAP condensates in log phase at pH 7.0, with a condensation value of 60.8 ± 2% compared to 43.2 ± 4% for WT cells (Fig. S4 A,B). To induce the stringent response, we added serine hydroxamate (100 μg/mL) which causes amino acid starvation. RNAP condensation decreased to 64.0 ± 2% in WT cells, but remained relatively high at 58.2 ± 3 % in Δ*relA* cells, despite nucleoid compaction (Fig. S4 A,B). These results are consistent with previous work implicating the stringent response in the dynamic redistribution of RNAP during amino acid starvation (5).

We next examined RNAP condensates in the Δ*relA* mutant during extreme acid stress. Since intracellular pH equilibrates 20-30 min after downshift to external pH 3.5 (Fig. 3C), we compared cells at t = 30 min. In contrast to WT, Δ*relA* mutants exhibit a significant decrease in RNAP condensation upon acid stress, from 66.5 ± 3% before to 61.8 ± 1% after pH downshift (Fig. 4C, D and S5A,B). This observation supports the hypothesis that the stringent response contributes to RNAP condensate stabilization during acid stress.

To distinguish the contributions of the two (p)ppGpp binding sites, we compared Δ*rpoZ* and Δ*dksA* mutants. Surprisingly, RNAP becomes even more dispersed during acid stress in Δ*rpoZ* mutants than in Δ*relA*, with condensation values from 68.52 ± 3% to 54.5 ± 2%, respectively (Fig. 4C, D and S7). In contrast, RNAP condensation in Δ*dksA* mutants does not change under acid stress, remaining at 72.1 ± 2% both before and after pH downshift(Fig. 4C, D and S5A,B). These results implicate the binding of (p)ppGpp to site 1 (at the β’ - ω interface), rather than site 2 (at the β’ - DksA interface), in the stabilization of RNAP condensates during extreme acid stress. We measured the total fluorescence intensity as a proxy for RNAP abundance across all conditions, and we show that decrease in condensation cannot be explained by a decrease in RNAP abundance (Fig S6 A,B,C).

To determine whether an *intracellular* pH downshift is necessary for the reduced condensation observed in the Δ*relA* and Δ*rpoZ* mutant backgrounds, we compared RNAP condensation in all strains after 30 min at external pH 7.0 or 5.5. We found that *ΔrelA, ΔrpoZ* and *ΔdksA* mutants have no RNAP condensation defect at pH 7.0 (Fig. 4E, F), nor growth defect (Fig. S7), showing that the stringent response is not necessary for RNAP condensation in optimal growth conditions. Moreover, RNAP condensation does not change during mild acid stress in any of the strains, confirming that these deletions do not affect condensate formation or stability at neutral cytoplasmic pH (Fig. 4E, F). These results support our hypothesis that the decrease in intracellular pH triggers the stabilization of RNAP condensates in WT cells, or their dissolution in *ΔrelA* and *ΔrpoZ* mutants.

The nucleolus is known to play a protective role for RNAs during various stresses (9, 83). Following the same logic, stable bacterial RNAP condensates may protect the transcriptional machinery during acute acid stress, allowing fast growth recovery, and as such contribute to cell fitness.

### Stabilization of RNAP condensates during acid stress correlates with survival

To determine whether stable RNAP condensates contribute to cell fitness during acid stress, we performed a colony forming unit (CFU) assay before and after 30 min of exposure to external pH 3.5 (see Materials and Methods) (Fig. 5A, shaded region). We then calculated the survival ratio, defined as the number of viable cells present after acid stress divided by the number of viable cells present before the stress (Fig. 5B). Only 4 ± 1% of WT cells survived acid stress, as reported previously (39). Unexpectedly, the survival ratio of the Δ*relA* mutant, 3 ± 1%, was similar to that of WT, despite the acid-induced decrease in RNAP condensation (Fig. 4D, Fig. 5B, yellow inset). Conversely, the Δ*rpoZ* mutant, which has even lower RNAP condensation during acid stress (Fig. 4D), and the Δ*dksA* mutant, which maintains a high level of condensation at external pH 3.5, each had much worse survival ratios: 0.07 ± 0.01% and 0.04 ± 0.03%, respectively (Fig. 5B, blue and red insets).

**Fig. 5.**
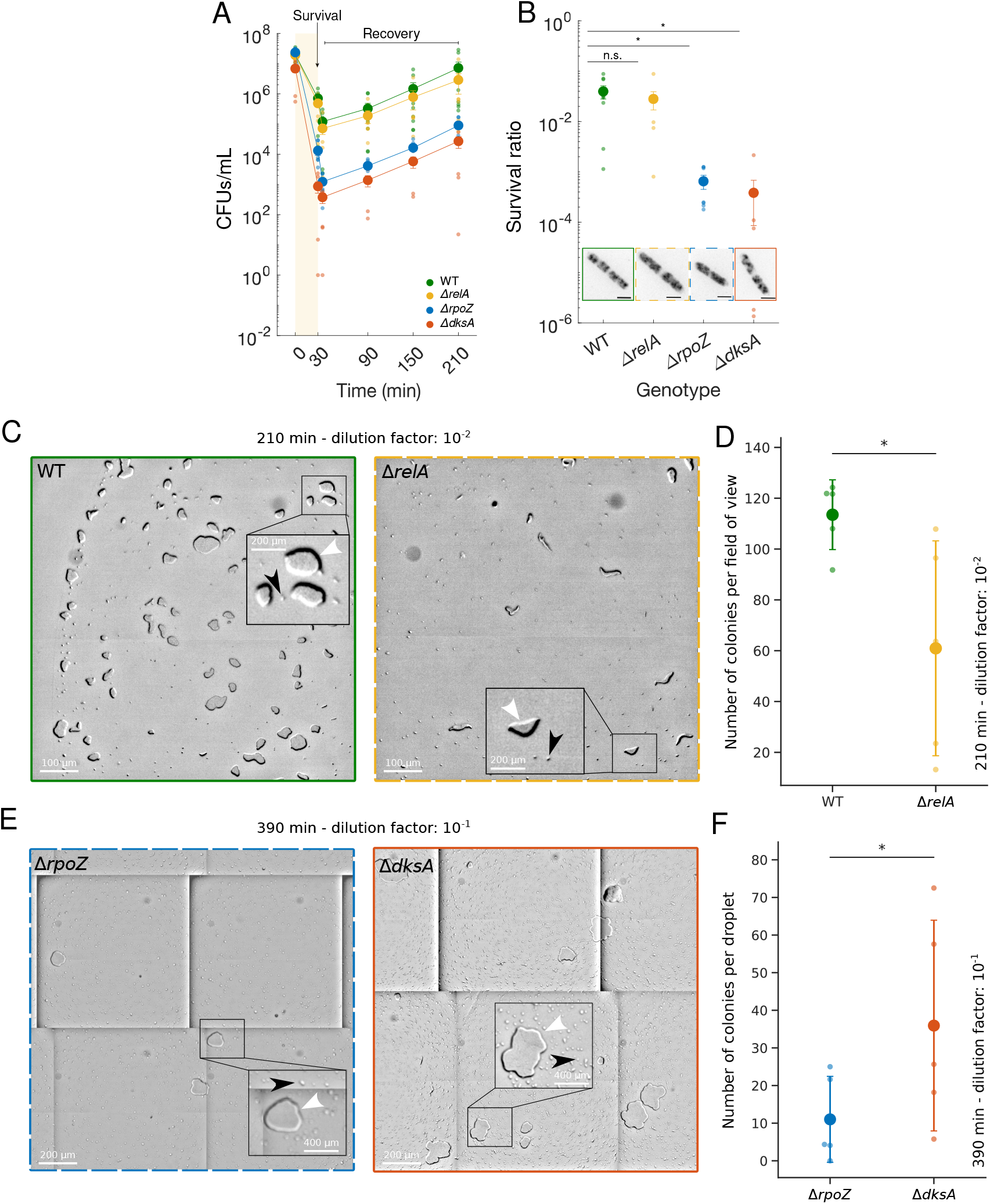
(A) Viable cell density of bulk liquid cultures of each genotype over time, before (0 min), during (30 min), and after recovery from downshift to pH 3.5. CFUs, colony forming units. (B) Survival ratio of each genotype after acid stress expressed as the fraction of viable cells at t = 30 min relative to t = 0 min. Inset: inverted fluorescence images of RpoC-mCherry in fixed cells at t = 30 min. Solid borders indicate genotypes that maintain RNAP condensates during acid stress; dashed borders indicate genotypes whose condensates dissolve. Scale bar, 2μm. (C) Representative images of colonies of WT and Δ*relA* during recovery from acid stress. White arrowheads indicate a single colony, black arrowheads indicate a single cell. Solid border indicates genotype that maintains RNAP condensates during acid stress; dashed border indicates genotype whose condensates dissolve. (D) Colony counts of WT and Δ*relA* cells subjected to pH 3.5 for 30 min, diluted 1:100, and recovered on agar plates for 3 hours. Values presented are the number of colonies per field of view, normalized OD_600_ at the time of plating. Transparent data points represent individual biological replicates; solid markers represent mean and standard error of N = 5 replicates for each genotype. p-values calculated using Kruskal-Wallis H test. *: p < 0.05. (E) Colony counts of Δ*rpoZ* and Δ*dksA* cells subjected to pH 3.5 for 30 min, diluted 1:10, and recovered on agar plates for 6 hours. Values presented are the total number of colonies observed normalized by the measured OD_600_. Transparent data points represent individual biological replicates; solid markers represent mean and standard error of N = 5 replicates. p-values calculated using Kruskal-Wallis H test. *: p < 0.05. (F) Representative images of Δ*rpoZ* and Δ*dksA* colinies during recovery from acid stress. White arrowheads indicate a single colony, black arrowheads indicates a single cell. Solid border indicates genotype that maintain RNAP condensates during acid stress; dashed border indicates genotypes whose condensates dissolve.

Our results were highly variable, especially for ΔdksA, which had the fewest CFUs at all time points (Fig. 5A). To probe survival and recovery from acid stress more directly, we visualized microcolony formation. Following 30 min at external pH 3.5, we collected samples of each strain; diluted them as needed; transferred 5 μL aliquots onto agar plates; incubated at 37°C for 3 or 6 hours; and acquired images with a high magnification (100x) large field stereo microscope (Axiozoom, Zeiss) (Fig. 5C, E). Finally, we counted the number of microcolonies and compared them within each survival group (i.e. high: WT and ΔrelA, and low: ΔrpoZ and ΔdksA) (Fig. 5D,F).

This approach allowed us to detect subtle differences in survival among our 4 bacterial strains. Consistent with the bulk CFU assays (Fig. 5A, B), we identified two groups based on the dilution factor and incubation time needed to observe individual microcolonies: 1) WT and Δ*relA* were diluted 100 times and observed after 3 hours of recovery from acid stress (Fig. 5C), and 2) Δ*dksA* and Δ*rpoZ* were diluted 10 times and observed after 6 hours of recovery from acid stress (Fig. 5E). That there are two distinct groups indicates that survival from acid stress is mediated primarily by the ω subunit and DksA, independently of RNAP condensation and (p)ppGpp. Furthermore, the greater sampling resolution of this microcolony assay allowed us to differentiate survival within each group. Indeed, WT had higher colony counts compared to Δ*relA* (Fig. 5D), and Δ*dksA* had higher colony counts compared to Δ*rpoZ* (Fig. 5F). This correlation between RNAP condensate maintenance and survival suggests a functional role for RNAP condensates during acid stress.

## Discussion

In this work, we show that bacterial RNAP condensates are sensitive to environmental physico-chemical cues. We propose that RNAP condensates formed during fast growth are stabilized during acid stress. In contrast to previous studies (4, 5), we find that fast cell growth is not required for the maintenance of RNAP condensates when cells are exposed to acid stress (Fig. 1). We also demonstrate that RNAP condensates become insensitive to hexanediol during acid stress(Fig. 2). Together, these results reveal a stress-induced stabilization of RNAP condensates which could reflect a transition and/or a change in the nature of interactions inside the condensates.

In addition, we show that the stringent response contributes to this condensate stabilization. Indeed, RNAP condensation decreases under acid stress in Δ*relA* and Δ*rpoZ* – but not Δ*dksA –* mutants, pointing to an important role for (p)ppGpp-binding site 1 in condensate stabilization (Fig. 4). This result is particularly surprising because the stringent response acts predominantly through (p)ppGpp-binding site 2 [22] during amino acid starvation, when it triggers dissolution of RNAP condensates (5). In contrast, mild acid stress, which does not induce intracellular pH downshift in WT (Fig. 3A, B), does not induce condensate dissolution in Δ*relA* and Δ*rpoZ* mutants (Fig. 4E, F) suggesting that RNAP condensates maintenance is specifically mediated by a low pH-induced stringent response.

Finally, we also demonstrate a weak correlation between RNAP condensate stabilization and cell fitness (Fig. 5). Although ω and DksA are the major contributors to fitness during acid stress, we nevertheless show that RNAP condensate maintenance increases survival relatively (Fig. 5). Indeed, Δ*dksA* mutants, which maintain their condensates during acid stress, have increased survival compared to Δ*rpoZ, which do not*. Similarly, WT has better survival than Δ*relA* mutants.

### Driving forces of RNAP condensate stabilization

RNAP condensates were first observed during fast growth (3, 4, 5), which highlighted their apparent link with transcription activity. During acid stress, however, we observe the persistence of RNAP condensates despite growth arrest. This result shows that RNAP condensates can be decorelated from growth and raises questions about the driving force(s) of condensate stabilization during acid stress. Intriguingly, our results show that acid stress increasingly impacts survival over time but we do not observe an associated shift in condensation which suggests that at this stage condensate maintenance is independent of bacterial activity. Weng and colleagues (3) found that the spatial organization of RNAP condensates depends not only on transcription but also on nucleoid compaction state. Indeed, reducing compaction by inhibiting gyrase activity leads to a partial dispersion of RNAP condensates. We show here that the nucleoid remains compact during acid stress (Fig. S3) and could as such contribute to the maintenance of RNAP condensates. This correlation is consistent with Weng.

Here we have only looked at condensates after 30 min of acid stress, even if a large proportion of cells are dead, we cannot exclude some adaptation mechanisms during early steps of acidification. In particular, ω and DksA are known to participate in gene expression regulation during nutritional stress. The differential behavior between the two mutants lacking one of these proteins could mean that ω contributes to gene expression adaptation with the requirement of (p)ppGpp during extreme acid stress, thus maintaining a high level of transcription, which in turn maintains RNAP condensates. Conversely, DksA would not contribute less to gene expression during extreme acid stress, and as such its deletion does not impact condensate maintenance. Better survival of Δ*dksA* compared to Δ*rpoZ* supports this hypothesis.

Another possible driving force for acid-induced stabilization of RNAP condensates is solidification of the cytoplasm. Indeed, acidification and low metabolic activity have been shown to impact the material state of the cytoplasm in various model organisms. For example, in budding yeast, the cytoplasm acidifies and undergoes a reversible liquid-to solid-like state transition within minutes of energy depletion. This transition limits the mobility of large exogenous particles, as well as endogenous organelles, DNA loci, and mRNA-protein complexes (40, 41). A similar phenomenon has also been observed in bacteria upon ATP depletion and other conditions of reduced metabolic activity (42, 70). Furthermore, in crowded cell lysates, pH can dramatically reduce the diffusion of cytoplasmic particles in a size-dependent manner: the diffusion of small particles is unaffected by low pH, whereas large particles diffuse more slowly at low pH than neutral pH (43).

Together, these *in vitro* and *in vivo* observations suggest that a downshift in intracellular pH could trigger a liquid-to-solid-like transition in the bacterial cytoplasm, and thereby kinetically trap RNAP molecules within condensates during acid stress. However, Δ*relA* and Δ*rpoZ* mutants lose their condensates during acid stress, indicating that RNAP and other condensate components are sufficiently mobile, and able to disperse by diffusing through the cytoplasm. The distinct behavior of RNAP condensates in WT (and Δ*dksA*) versus the Δ*relA* and Δ*rpoZ* mutants suggests that RNAP condensate stabilization occurs through a local mechanism (i.e. at the level of intermolecular interactions within the condensate or with the DNA), rather than through a global one (i.e. physical state of the cytoplasm).

### Physico-chemical nature of RNAP condensates

Low pH results in protonation of amino acid side chains which can strongly modulate electrostatic interactions within or between molecules. In eukaryotic and prokaryotic organisms, the distribution of protein pIs is often bimodal (46) (Fig. S8). The two peaks are roughly centered around pH 6.0 and 9.0, which means that at intracellular pH below 4.0 (Fig. 3A), most proteins have a positive net charge inside the cytoplasm, as observed for β and β’ (Fig. 3). Moreover, *in vitro* (33, 37) and theoretical (35, 47) works have shown that deviation of the pH away from a protein’s pI is unfavorable for phase separation in cases where condensate formation is driven by hydrophobic interactions (31) due to both electrostatic repulsion between residues of the same charge sign and changes in protein solubility. These results are supported by observations in human and mouse skin where differentiating keratinocytes lose their liquid-like keratohyalin granules containing filaggrin (pI = 9.1) after intracellular pH downshift (16). Nevertheless, in *E. coli*, we find that native RNAP condensates do not dissolve during acid stress despite the intracellular pH dropping below the pIs of condensate components.

Ionic strength determines the relative strength of electrostatic interactions, and can therefore tune the driving forces underlying phase separation. At low salt concentration (high ionic strength), phase separation can be driven by electrostatic and/or hydrophobic interactions. With increasing salt concentrations (lowering ionic strength), the screening of electrostatic charges can cause condensates to dissolve, and ultimately induce a reentrant phase transition, in which condensates reform through hydrophobic and non-ionic interactions (48). Reentrant phase separation was also theoretically observed as a function of pH. In this case, reentrant behavior is driven by switching from hydrophobic to electrostatic interactions, and electrostatic to hydrophobic (35). Similarly, we speculate that pH-dependent modulation of the intermolecular interactions underlying RNAP condensates contributes to the differences in condensate stability at pH 7.0 and pH 3.5.

Oppositely charged molecules, together, can undergo phase separation, a phenomenon known as complex coacervation (49, 50). For example, DNA and RNA undergo complex coacervation with several different proteins like Zein and Tau, respectively (52,53,71). During acid stress, positively-charged RNAP is in close proximity to DNA and RNA, which are negatively charged. In addition, our results show that, while RNAP condensates are sensitive to hexanediol at intracellular pH 7.0, they become insensitive during acid stress. Hexanediol insensitivity is also observed for Tau, when undergoing phase separation through complex coacervation with RNA (72). Altogether, these observations suggest a change in the nature of intermolecular interactions inside RNAP condensates (Fig. 2C,D), from hydrophobic interactions at pH 7.0 to electrostatic interactions at pH 3.5.

### Stress-specific stringent responses

Divergence regarding the dynamics of phase separated bodies were observed in human cells (8) and in yeast (17) as a function of stressors. In *E. coli*, RNAP condensates rapidly dissolve upon nutrient starvation (5) (Fig. S4), but are stabilized during acid stress (Fig. 1). We thus show that the nature of stress can affect condensate properties in different and even opposite manner in bacteria, just like in eukaryotic cells. Further investigations are needed to understand the biological function of such stress-specific behavior, and why they are conserved across distinct domains of life.

Both phenotypes observed in bacteria, i.e. starvation-induced dissolution (55) and acid-induced stabilization (Fig. 4), are dependent on the (p)ppGpp synthase RelA. This shows that bacteria can actively regulate RNAP condensates during stress, and that passive, physicochemical mechanisms are not the only ones involved. In addition, the differential behavior of RNAP condensates to various stressors demonstrates that the stringent response is stress-specific and can trigger distinct downstream effects on bacterial cell organization. Much of the literature dissecting “the” stringent response has relied on nutrient starvation, specifically amino acid starvation, but our results underscore the importance of investigating additional stress conditions, including heat shock (74), phosphate starvation (75), DNA damage (76), glucose phosphate stress (77) and oxidative stress (78).

RNAP regulation by (p)ppGpp has mostly been studied from the point of view of DksA. Indeed, binding of (p)ppGpp to site 2 has a stronger effect on gene expression during amino acid starvation compared to site 1 (22) despite having similar affinities for each site (56). Furthermore, in ω depleted strains, DksA overexpression fully rescues RNAP response to (p)ppGpp (57). These observations seemingly diminish the importance of ω but we identify here a novel role for the ω subunit, distinct from DksA, in maintaining RNAP condensates during acid stress.

### Condensate stabilization and cellular fitness

Unexpectedly, RNAP condensates do not contribute to fitness during acid stress as directly as we could have thought. Indeed, ω and DksA seem to be the biggest contributors to cell fitness during acid stress, independently of RNAP condensation. In addition, Δ*relA* high survival indicates that omega and DksA activity is independent of the stringent response in contrast to what is observed during nutrient starvation, where (p)ppGpp increases ω and DksA effect (22). Interestingly, direct exposure of DksA to pH 6.0 *in vitro* induces conformational changes increasing its affinity to RNAP, rendering its effect on gene regulation as strong as DksA bound to (p)ppGpp (60). Similarly, in *E. coli*, the activity of the acid stress response factor lysine decarboxylase (LdcI) is independent of (p)ppGpp only at low pH (15) which, together with our results, suggests that, like LdcI, DksA becomes a (p)ppGpp independent actor of the stress response at low pH. Similar hypotheses could be developed regarding the ω subunit. In addition, we highlight an essential role for DksA in acid stress adaptation during the exponential phase, as previously observed during stationary phase (39, 60). Finally, mutations at the interface of β’ and ω arise during cellular acid adaptation experiments (61) strongly pointing to a particularly important role for ω during acid stress. How these mutations affect the acid-induced stabilization of RNAP condensates is an open direction for future investigations.

Nevertheless, we show that for similar survival ranges, namely survival of WT and Δ*relA* on the one hand, Δ*rpoZ* and Δ*dksA* on the other hand, maintenance of RNAP condensates favors survival. This result illustrates that, even if this effect is partially masked by the preponderant role of ω and DksA during acid stress, RNAP condensate maintenance is a viable stress adaptation strategy in bacteria, similarly to what is observed for the nucleoli in eukaryotes.

### Outlook

The functional role of RNAP condensates during fast growth at neutral pH remains to be fully explored. Their stabilization during acid stress seems to confer protection to RNAP against drugs, including antibiotics, and they do contribute to cellular fitness during or after acid stress. We show that ω and (p)ppGpp binding site 1 are clearly involved in the stabilization of RNAP condensates. Previously, ω has been shown to have a weaker effect on (p)ppGpp-dependent gene regulation compared to DksA (22), but in contrast, here we demonstrate that it has a role at the supramolecular level that DksA lacks. In our work, we highlight this phenomenon at extreme pH values, but several environmental contexts such as ATP depletion in bacteria (44), entry into dormancy and heat stress in yeast (40, 45, 68), are linked to downshifts in intracellular pH. It is thus necessary to investigate further how pH variations affect RNAP organization in other contexts, with and without ω, and how it can be linked to stress resistance or adaptation. Ultimately, our work opens up new perspectives regarding the function of ω, the RNA polymerase subunit whose role has remained a mystery since its discovery (61).

## Materials & Methods

### Strains and plasmids

Strains used in this study are listed in Table S1. P1 transduction was used to generate new strains (https://currentprotocols.onlinelibrary.wiley.com/doi/epdf/10.1002/0471142727.mb0117s79). Specifically, WLBS111 was generated by P1 transduction from JW0141-1 into WLBS100; WLBS121 was generated by P1 transduction from JW3624-1 into WLBS100; and WLBS122 was generated by P1 transduction from JW2755-3 into WLBS100. Plasmid pSCM001 was used to transform cells with SEpHluorin fusion protein (36).

### Growth conditions

Liquid cultures were inoculated from a single colony and grown overnight at 37°C in LB. Saturated cultures were diluted into fresh LB (pH 7.0) to an OD_600_ of 0.05, grown for another 16 h, and then diluted again to start each experiment. Cultures were grown at 37°C in a water-bath shaking incubator for the time of the experiment.

### Acid stress and time-course imaging

Cells were grown for 90 min in 25 mL flasks of fresh LB, then HCl (1 M) was added to bring the pH of the culture to the desired pH. To determine the volume needed, HCl was gradually added to 25 mL of clean LB while monitoring the pH with a pH-meter. When the target pH was reached, the total volume of HCl added was recorded (600 μL for pH 3.5 and 150 μL for pH 5.5). Cells were incubated for 30 more minutes in the acidified LB. Samples were then collected every 60 min; fixed in 4% formaldehyde with constant mixing for 30 min at 37°; washed three times in minimal M9 medium at room temperature; and stored at 4°C. pH was measured again after experiment to confirm cells were exposed to the correct value. Fixed cells were mounted on 1% M9-agarose pads and imaged on an inverted Leica DMI 6000B equipped with a 100X 1.46 NA objective lens, a spinning disk confocal head (Yokogawa CSU10) and an EM-CCD camera (Hamamatsu ImagEM). Five to ten fields of view were randomly acquired for each time point.

### Image analysis

Image analysis was done using custom Java/ImageJ/Python scripts. Cells were segmented from bright field images and fluorescence intensity values of all pixels within a cell were extracted. Condensation was calculated as described in (4,63). Briefly, β’-mCherry signal intensity (I) in cells was normalized such that *I*_*n*_ = (*I* − *I*_*min*_)/(*I*_*max*_ − *I*_*min*_). By measuring the percent of intensity values satisfying the condition *I*_*n*_ *<* 0.5, we obtain a single condensation value per cell, corresponding to the proportion of the background signal in the cell. This value increases upon protein condensation and allows comparison across different conditions.

### Drug treatments

#### Hexanediol

Following 30 min acid stress, cultures were treated with hexanediol to reach a final concentration of 5%. Each condition had 3 biological replicates, performed on different days to account for variation in individual experiments. Sampling occurred moments prior to and 5 min after drug addition. Samples were fixed and imaged as described above.

#### Serine hydroxamate

Cultures were treated with Serine Hydroxamate to reach a final concentration of 100 μg/mL. Each condition had 3 biological replicates, performed on different days to account for variation in individual experiments. Sampling occurred moments prior to and 20 min after drug addition. Samples were fixed and imaged as described above.

### Molecular structure of the RNAP complex

Molecular structure #5VSW (64) containing RNAP complex as well as DksA and (p)ppGpp was selected on RCSB Protein Data Bank website (https://www.rcsb.org/). Electrostatic potential maps of the complex were calculated at each pH on the APBS-PDB2PQR software suite web server (https://server.poissonboltzmann.org/) (65). UCSF ChimeraX (https://www.rbvi.ucsf.edu/chimerax/) (66) was then used to map the calculated electrostatic potential surface onto the molecular structure. UCSF ChimeraX is developed by the Resource for Biocomputing, Visualization, and Informatics at the University of California, San Francisco, with support from National Institutes of Health R01-GM129325 and the Office of Cyber Infrastructure and Computational Biology, National Institute of Allergy and Infectious Diseases.

### Survival and recovery assays

#### Spot assay

To measure survival, cultures were sampled before (time 0 min), 30 min and 90 min after pH downshift. Serial dilution was done for each sample and 5 μL was deposited on agar plates. Plates were incubated at 37°C for 24 hours and the number of colonies was counted manually on the lowest density dilution. Survival was calculated as the ratio of viable cells during acid stress to that before stress.

#### CFU assay

To calculate survival, cultures were sampled just before (time 0 min) and 30 min after acid stress. The number of viable cells in each sample (CFU/mL) was determined by serial dilution and the pour plate technique. Briefly, 100 μL of diluted samples were added to empty plates, then molten LB-agar (cooled to 45°C, containing chloramphenicol) was poured into the plates. Plates were gently rotated in a circular motion to achieve uniform distribution of samples and left at room temperature until the agar solidified. Plates were then incubated at 37 °C for 48 hours and the number of colonies was counted manually. Survival was calculated as the ratio of viable cells after acid stress to that before stress.

#### Microcolony assay

To monitor micro colony formation on agar plates, cultures were sampled 30 min after acid stress. Samples were diluted by a factor of 100 or 10 and then 5 μL were deposited on agar plates. Plates were incubated at 37°C for 3 or 6 hours respective to their dilution factor and pictures of the whole deposited area were acquired using a stereoscopic microscope at 100x magnification (Zeiss, Axiozoom). The number of colonies per field of view or the total number of colonies was counted manually. The number of colonies was normalized by the measured OD_600_ at time 0 min.

## Acknowledgements

We thank members of the Weber lab for helpful discussions. Deletion strains from the Keio collection were obtained from the Coli Genetic Stock Center, which is supported by the National Science F (DBI-0742708). This work was supported by the Natural Sciences and Engineering Research Council of Canada RGPIN-2017-04435 (to S.C.W.). This research was undertaken, in part, thanks to funding from the Canada Research Chairs Program (CRC-2020-00325 to SCW).

