## Supplementary material for "Omega stabilizes RNA polymerase condensates and contributes to cellular fitness during acid stress": Supllemental Information

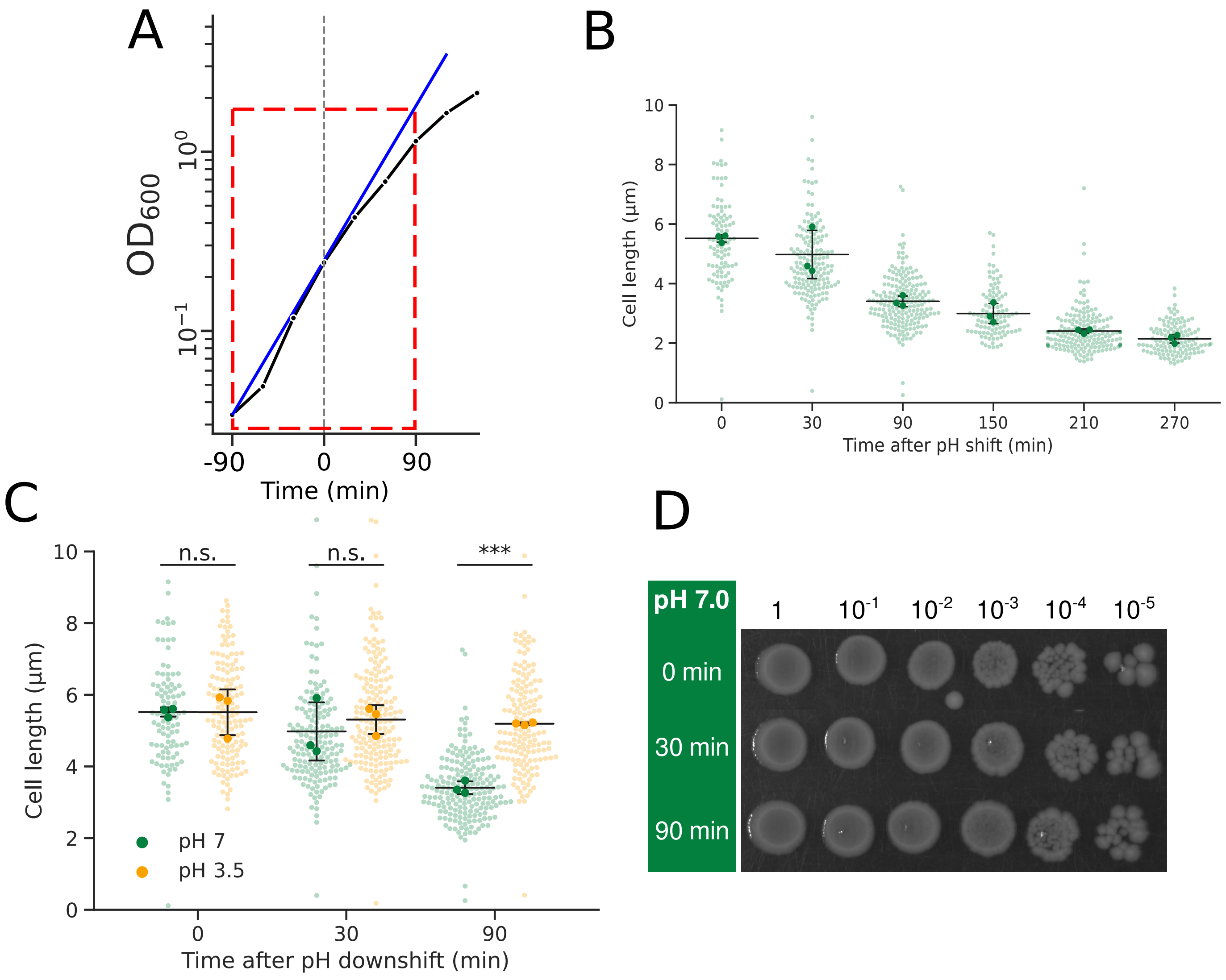


**Fig. S1** – (A) Semi-log representation of the growth curve shown in Fig.1A. Blue line represents growth rate in exponential phase. (B) Length of *E. coli* cells over time after entry into exponential phase at pH 7 (0 min: n = 98, 30 min: n = 168, 90 min: n = 191, 150 min: n = 190, 210 min: n = 294, 270 min: n = 211). Transparent data points correspond to individual cells; solid markers represent the population mean of each biological replicate; black bars represent mean and standard error of N = 3 replicates. (C) Length of *E. coli* cells over time after pH downshift (pH 7.0 - 0 min: n = 98, 30 min: n = 168, 90 min: n = 191; pH 3.5 - 0 min: n = 146, 30 min: n = 238, 90 min: n = 239). WT data are repeated from panel B. Transparent data points correspond to individual cells; solid markers represent the population mean of each biological replicate; black bars represent mean and standard error of N = 3 replicates. p-values calculated by t-test. n.s.: non-significant; ***: $p < 0.001$. (D) Survival assay at various times after entry into exponential phase at pH 7.0. Each column represents a dilution of 5 µL droplet. Experiment was repeated 3 times.

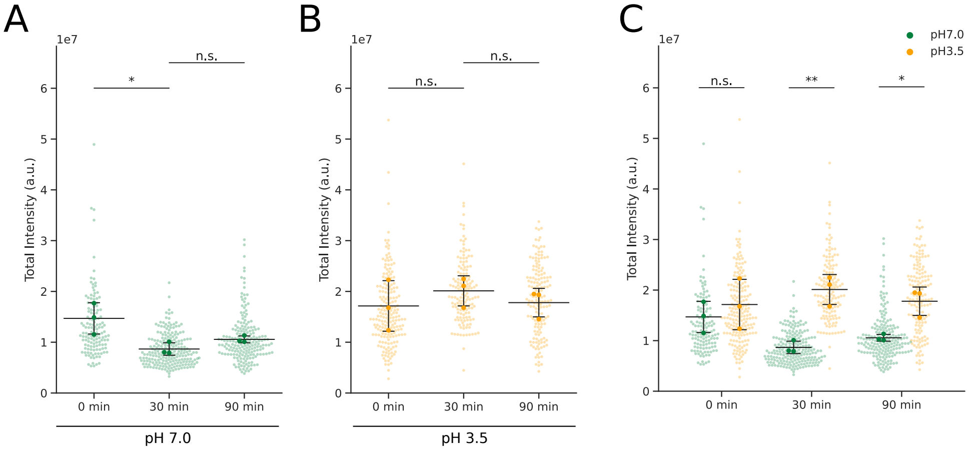


**Fig. S2** – Total fluorescence intensity of RpoC-mCherry per cell at various time points after pH downshift at (A) pH 7.0 (0 min: n = 135, 30 min: n = 216, 90 min: n = 197) and (B) pH 3.5 (0 min: n = 194, 30 min: n = 148, 90 min: n = 157). Transparent data points correspond to individual cells; solid markers represent the population mean of each biological replicate; black bars represent mean and standard error of N = 3 replicates. p-values calculated by ANOVA test. n.s.: non-significant; *: $p < 0.05$.

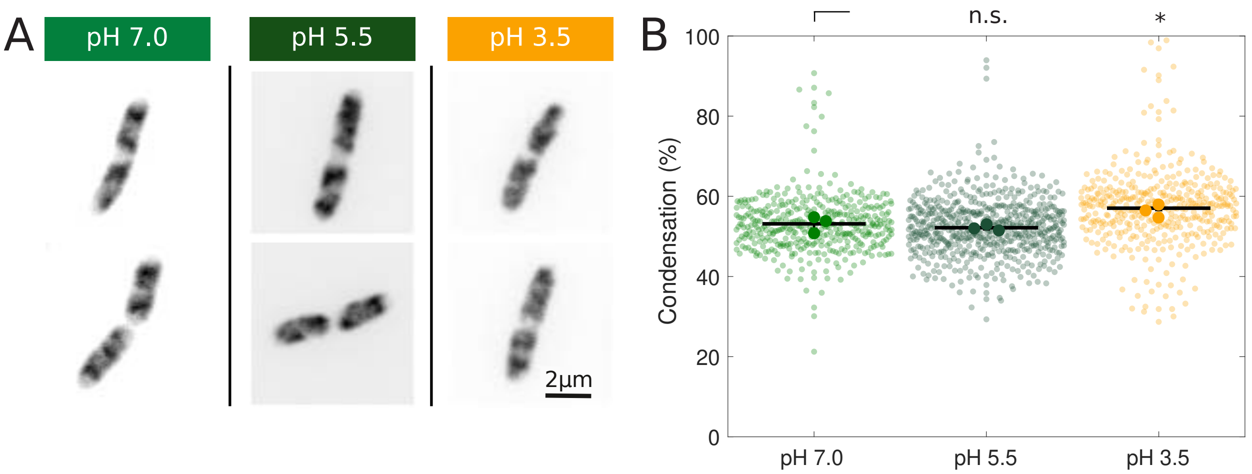


**Fig. S3** – (A) Fluorescence images (contrast inverted) of fixed cells expressing HupA-mCherry shifted to pH 7.0, pH 5.5, or pH 3.5 for 30 min. (B) Condensation of the nucleoid (HupA-mCherry) before and after pH downshift. (pH 7.0 - 0 min: n = , 30 min: n = ; pH 5.5 - 0 min: n = , 30 min: n = ; pH 3.5 - 0 min: n = , 30 min: n = ). Transparent data points correspond to individual cells; solid markers represent the population mean of each biological replicate; black bars represent mean and standard error of N = 3 replicates. p-values calculated by ANOVA

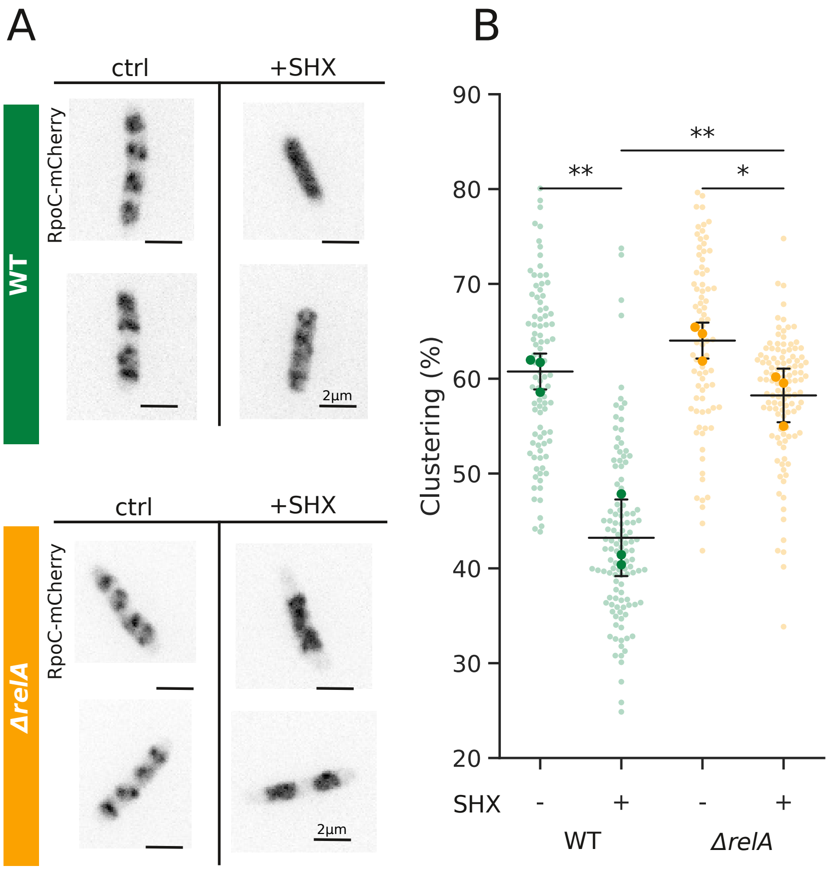


**Fig. S4** – RNAP condensates of cells treated with serine hydroxamate (SHX) to mimic amino acid starvation and induce the stringent response. (A) Fluorescence images (contrast inverted) of fixed cells expressing RpoC-mCherry in WT (top) and Δ*relA* (bottom) strains after 20 min SHX treatment. (B) Condensation of RpoC-mCherry before (-) and after (+) SHX treatment in WT (+: n = 92; -: n = 118) and Δ*relA* (+: n = 80; -: n = 114) cells. Transparent data points correspond to individual cells; solid markers represent the population mean of each biological replicate; black bars represent mean and standard error of N = 3 replicates. p-values calculated by ANOVA test. *: $p < 0.05$: **: $p < 0.01$.

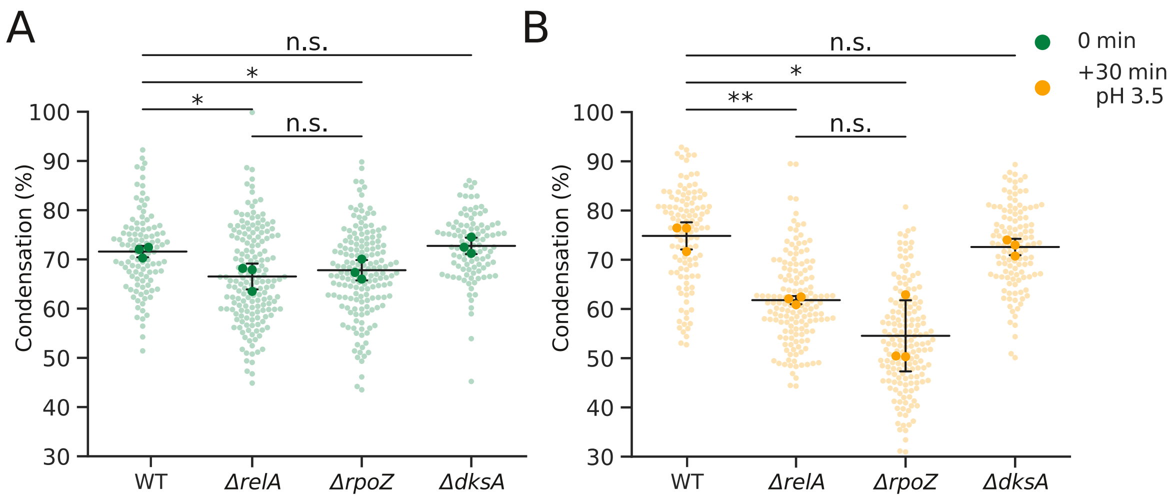


**Fig. S5** – Condensation of RpoC-mCherry before (A) and after (B) pH downshift across different strains: WT (pH 7.0: n = 136, pH 3.5: n = 52), Δ*relA* (pH 7.0: n = 137, pH 3.5: n = 130), Δ*rpoZ* (pH 7.0: n = 98, pH 3.5: n = 233) and Δ*dksA* (pH 7.0: n = 148, pH 3.5: n = 110) before (A) and after (B) pH downshift. Same data as Fig. 4D but rearranged for comparison. Transparent data points correspond to individual cells; solid markers represent the population mean of each biological replicate; black bars represent mean and standard error of N = 3 replicates. p-values calculated by ANOVA test. n.s.: non-significant; *: $p < 0.05$: **: $p < 0.01$.

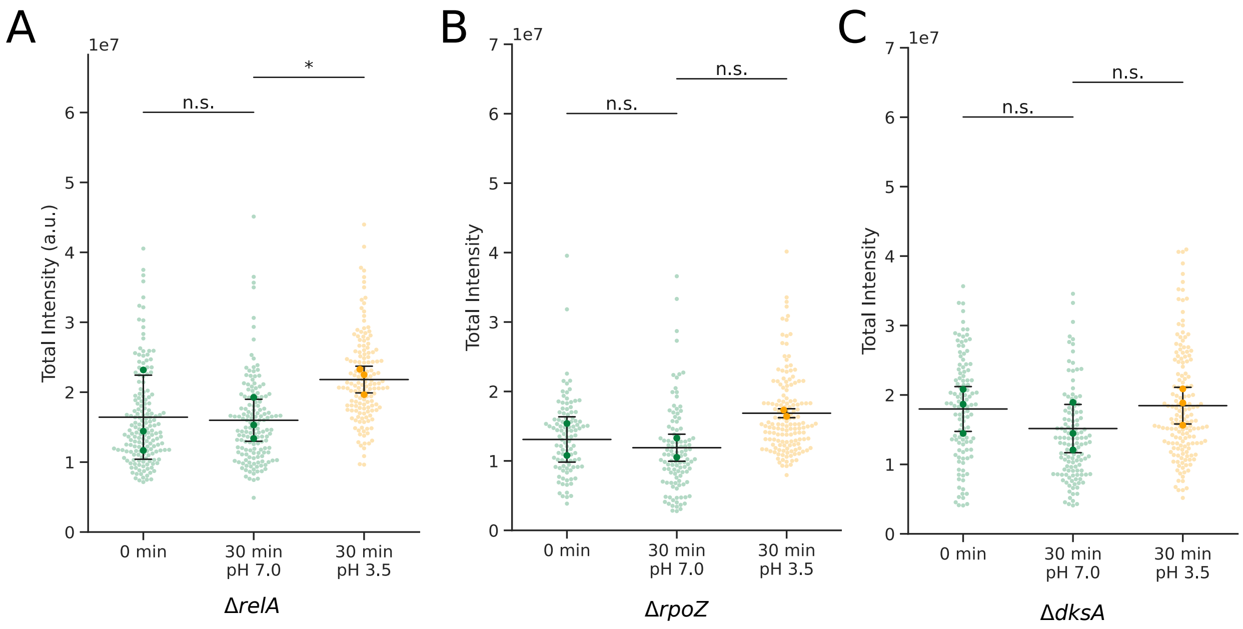


**Fig. S6** – Total fluorescence intensity of RpoC-mCherry per cell in mutant strains before (0 min) or after (30 min) pH downshift to pH 7 or pH 3.5. (A) Δ*relA* (0 min: n = 171, 30 min pH 7.0: n = 137, 30 min pH 3.5: n = 149), (B) Δ*rpoZ* (0 min: n = 164, 30 min pH 7.0: n = 98, 30 min pH 3.5: n = 166) and (C) Δ*dksA* (0 min: n = 103, 30 min pH 7.0: n = 110, 30 min pH 3.5: n = 121). Transparent data points correspond to individual cells; solid markers represent the population mean of each biological replicate; black bars represent mean and standard error of N = 3 replicates. p-values calculated by ANOVA test. n.s.: non-significant.

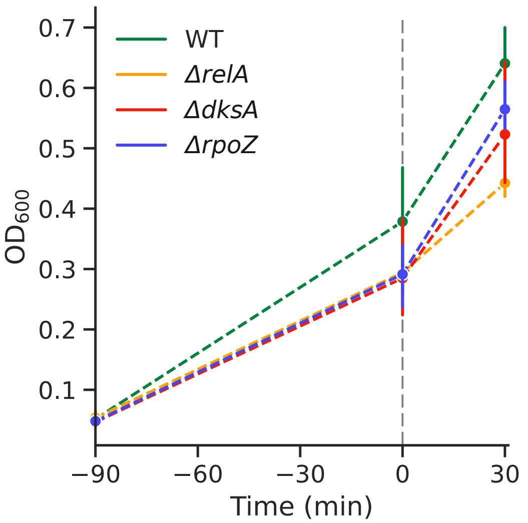


**Fig. S7** – OD600 of cultures of WT, Δ*relA*, Δ*rpoZ* and Δ*dksA* growing at pH 7.0 in rich media (LB). Solid markers represent mean of N = 3 biological replicates; bars represent standard error of replicates

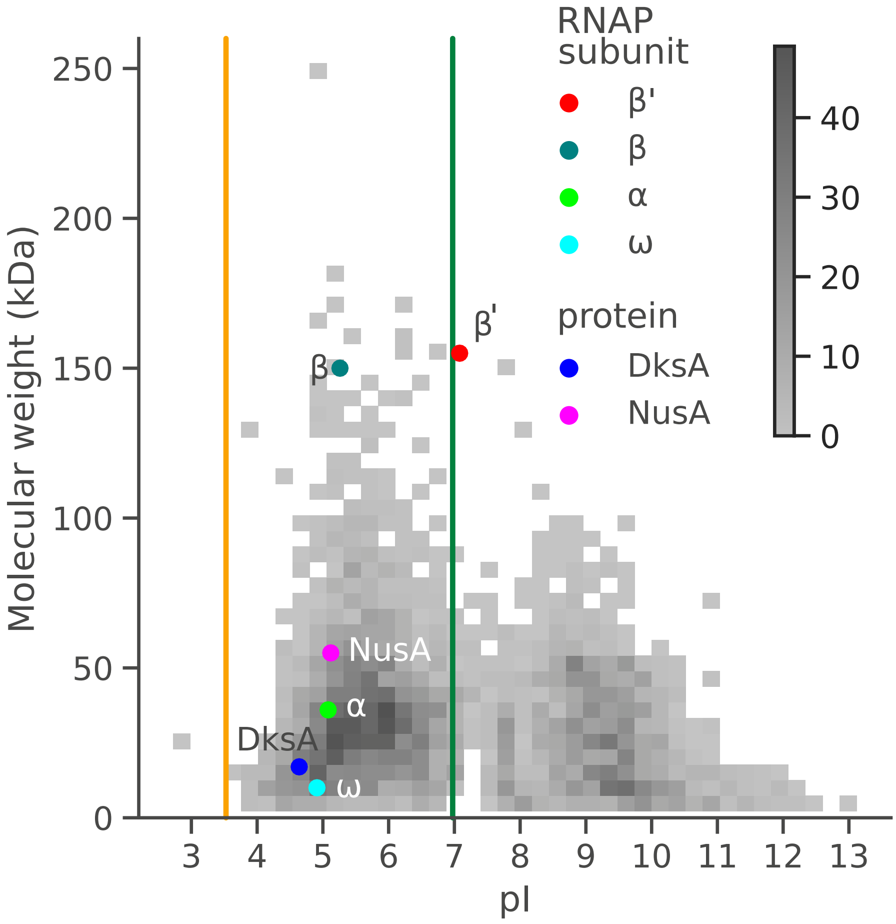


**Fig. S8** – Distribution of isoelectric points (pI) across *E. coli* proteome. Vertical lines represent intracellular pH for cells shifted to pH 7.0 (green) or pH 3.5 (yellow). Proteins of interest are represented by solid markers. Color scale represents the number of proteins per square.

| Strain | Genotype | Source |
|---|---|---|
| MG1655 | F-, λ-, *rph-1* | CGSC |
| JW0141-1 | BW25113; Δ*dksA-761::kan* | CGSC |
| JW3624-1 | BW25113; Δ*rpoZ-754::kan* | CGSC |
| JW2755-3 | BW25113; Δ*relA-782::kan* | CGSC |
| RRL265 | AB1157; *rpoC-mCherry frt-cat* | Ladouceur *et. al*, PNAS, 2020 |
| WLBS100 | MG1655; *rpoC-mCherry frt-cat* | Ladouceur *et. al*, PNAS, 2020 |
| WLBS111 | MG1655; *rpoC-mCherry,* Δ*dksA-761::frt-kan* | This study |
| WLBS121 | MG1655; *rpoC-mCherry,* Δ*rpoZ-754::frt-kan* | This study |
| WLBS122 | MG1655; *rpoC-mCherry,* Δ*relA-782::frt-kan* | This study |

**Table S1** – Bacterial strains used in this study. "CGSC" stands for "Coli Genetic Stock Center". New strains were constructed using P1 transduction protocol from Thomason LC, Costantino N, Court DL (2007) E. coli Genome Manipulation by P1 Transduction. Current Protocols in Molecular Biology, 79(1), 1.17.1–1.17.8. doi: 10.1002/ 0471142727.mb0117s79. WLBS111: P1 transduction from JW0141-1 into WLBS100 - WLBS121: P1 transduction from JW3624-1 into WLBS100 - WLBS122: P1 transduction from JW2755-3 into WLBS100
